# Connectivity dynamics and cognitive variability during aging

**DOI:** 10.1101/2022.01.26.477817

**Authors:** G. Jauny, F. Eustache, T. Hinault

## Abstract

Aging is associated by cognitive changes, with strong variations across individuals. One way to characterize this individual variability is to use techniques such as magnetoencephalography (MEG) to measure the dynamics of neural synchronization between brain regions, and the variability of this connectivity over time. Indeed, few studies have focused on fluctuations in the dynamics of brain networks over time and their evolution with age. We therefore characterize aging effects on MEG phase synchrony in healthy young and older adults from the Cam-CAN database. Age-related changes were observed, with an increase in the variability of brain synchronization, as well as a reversal of the direction of information transfer in the default mode network (DMN), in the delta frequency band. These changes in functional connectivity were associated with cognitive decline. Results suggest that advancing age is accompanied by a functional disorganization of dynamic networks, with a loss of communication stability and a decrease in the information transmitted. This could be partly due to the loss of integrity of the network structure.

## Introduction

With an increasing number of people over 65, the world population is aging. Aging is associated with a reduced efficiency of cognitive functioning, that primarily affects memory and executive processes (e.g., Hedden & Gabrieli, 2004). However, some individuals show a major decline while others maintain cognitive performance similar to young adults (e.g., Hultsch *et al*., 2008). Recent research aims to better understand these individual differences during aging. Such variability across individuals has been associated with concepts of maintenance and cognitive reserve (Cabeza *et al*., 2018; Stern *et al*., 2020). Maintenance (Nyberg et al., 2012) corresponds to the preservation of similar cognitive and brain functioning to that of younger individuals with advancing age, while cognitive reserve corresponds to compensatory functional adjustments associated with the preservation of cognitive performance in the presence of structural changes. Cognitive reserve and maintenance can account for individual differences in aging-and pathology-related effects, and have been extensively investigated at both structural and functional levels (e.g., Stern *et al*., 2020). However, the contribution of the temporal dynamics of brain communications underlying cognitive reserve remains under-investigated. As changes in brain dynamics are expected to occur long before the disconnection associated with atrophy and brain lesions, this could yield highly sensitive elements on individual differences with age.

Neuroimaging research in healthy aging has been primarily conducted using functional methods with high spatial resolution (e.g., positron emission tomography (PET) or functional MRI (fMRI)). These methods have provided insights into the anatomical and functional changes that occur with age, including changes in brain activity (Cabral *et al*., 2017; Smitha *et al*., 2017). These techniques also enable the study of brain connectivity changes. Connectivity measures are sensitive to cognitive changes and differences between individuals (e.g., Hedden *et al*., 2016). Studies showed, for example, that the cognitive decline observed in normal aging may be due to functional connectivity disruptions, particularly in the default-mode network (DMN; this network is mainly activated when no task is requested from the participant; Andrews-Hanna *et al*., 2007). The concept of cognitive reserve itself has also emerged in part from fMRI studies, as individuals with a higher cognitive reserve showed fewer brain and cognitive alterations than individuals with a lower level of cognitive reserve (Stern, 2009). The contribution of these methods in the precise localization of brain activity and in the study of brain networks is therefore undeniable. However, due to their constrained temporal resolution, age-related changes on the dynamics of the networks involved remain largely understudied. The use of methods with high temporal resolutions, such as magnetoencephalography (MEG) and electroencephalography (EEG; e.g., Baillet, 2017), can provide sensitive and specific elements on individual differences associated with cognitive aging.

Brain activity is characterized by its spectral complexity, and can be distinguished according to its dominant frequency (delta, theta, alpha, beta, gamma). The delta waves (1-3Hz) are the slowest, while the gamma waves (40+Hz) are the fastest. Previous work highlighted that these brain rhythms are associated with different cognitive functions (Buszaki *et al*., 2006), for example the gamma frequency band is associated with information processing in higher-order cognitive tasks. Previous MEG studies show that networks activated at rest are activated periodically and in different frequency bands (de Pasquale *et al*., 2010). A decrease in functional connectivity has been observed during aging (Wig, 2017). Moreover, previous work has shown that in older individuals, activations and couplings are reduced in the alpha and gamma frequency bands, and increased in the delta frequency band (Vlahou, 2014). This slowing of neural activity (Celesia, 1986) has been linked to decreased cognitive performance (Toth et al., 2014), and slower information processing speed (Anderson & Craik, 2017). Conversely, the preservation of this neural activity allows cognitive abilities to be maintained with age. However, previous M/EEG studies have mainly focused on the average of activations and connectivity over long periods of time (see Courtney & Hinault., 2021, for a review), and therefore do not provide insight into the dynamics of brain activities or their association with cognitive changes. It is therefore important to study the fluctuations of brain communications over time.

Spontaneous fluctuations of brain activity have long been considered as noise to be eliminated and/or controlled for. They are now considered as a fundamental aspect of brain communications (e.g., Uddin, 2020). Recent work has demonstrated the importance of sustained synchronization between brain regions for performance in complex cognitive tasks (e.g., Daume *et al*., 2017). Moreover, disrupted synchronization has been associated with cognitive decline with age (Hinault *et al*., 2020). However, fluctuations of activity have not been considered in light of individual differences during aging. Impaired stability of brain network dynamics could lead to the neurocognitive changes observed with advancing age (Voytek & Knight, 2015). The directionality of connectivity between neuronal oscillations may also play a role in the transmission of neuronal communications. Therefore, changes in dynamic connectivity would take place in order to maintain cognitive performance, while failure to make these changes would lead to cognitive decline (Ariza *et al*., 2015).

Here, we investigated the stability and variability of resting brain networks’ synchrony over time in young and older healthy participants from the Cam-CAN (*Cambridge Centre for Ageing and Neuroscience*) database (e.g., Shafto *et al*., 2014; Taylor *et al*., 2017). This database includes multimodal neuroimaging data (MEG, f/MRI) as well as cognitive performance assessment in each individual. Analyses were focused on the four main resting-state networks found activated at rest that are the (default network, salience network, left and right fronto-parietal networks). Our objectives were twofold: i) To study changes in dynamic connectivity with age: Between young and old individuals, we hypothesized differences in functional networks, as well as greater variability in the activity of these networks; ii) To investigate the relationships between changes in dynamic connectivity and cognitive changes: We expected that stability in synchronization and directionality of connectivity over time would be associated with better cognitive performance with age, compared to high variability in these measures. Preservation of this neural activity would help maintain cognitive abilities with age.

## Methods

### Participants

We analysed data from 46 young (29 women and 17 men; aged 22-29 years) and 46 older healthy adults (29 women and 17 men; aged 60-69 years; see participant demographics characteristic in Table 1). Participants were selected from the Cam-CAN database (e.g., Shafto *et al*., 2014; Taylor *et al*., 2017), in line with demographic characteristics of individuals recruited in previous work (e.g., Coquelet *et al*., 2017; Hinault *et al*., 2020). All participants were right-handed, showed normal cognitive functioning (Montreal Cognitive Assessment (MoCA) score >26; Nasreddine. *et al*., 2005), and no neurological or psychiatric condition.

**Table 1:** Demographics and scores for both groups younger and older participants

| Variables | Young adults | Older adults | p-value |
| --- | --- | --- | --- |
| Number of participants | 46 | 46 | - |
| Number of females | 29 | 29 | - |
| Age | 26.5 | 64.5 | - |
| **Years of education** | **22.2** (2.873) | **19.1** (3.262) | **0.001** |
| **MMSE** | **29.5** (0.863) | **28.9** (1.173) | **0.013** |
| **VSTM** | **0.5** (0.088) | **0.4** (0.069) | **0.001** |
| **Cattell** | **37.8** (3.628) | **30.5** (6.285) | **0.001** |
| **Hotel_Num_rows** | **4.7** (0.585) | **4.3** (1.008) | **0.018** |
| **Hotel_Time** | **227.7** (119.796) | **326.9** (194.305) | **0.005** |

### Behavioural measures

A detailed description of the behavioural measures can be found in supplementary materials and in Shafto *et al*. (2014) and Taylor *et al*. (2017). Cognitive performance was assessed with the Mini-Mental State Evaluation (MMSE; Folstein *et al*., 1975) used as a measure of general cognitive functioning, the Visual Short-Term Memory (VSTM; Vogel et al., 2001) which measures working memory, the Cattell test (Horn & Cattell, 1966) which is a measure of reasoning ability and the Hotel Test (Shallice & Burgess, 1991) which assesses planning abilities.

### MEG and structural MRI data acquisition

Resting brain activity was measured for 10 minutes (sampling rate: 1kHz, bandpass filter: 0.03-330 Hz) with a 306-channel MEG system. Participants’ 3D-T1 MRI images were acquired on a 32-channel 3T MRI scanner. The following parameters were used: repetition time = 2250 ms; echo time = 2.99 ms; inversion time = 900 ms; flip angle = 9 degrees; field of view = 256 mm x 240 mm x 192 mm; voxel size = 1 mm; GRAPPA acceleration factor = 2; acquisition time = 4 minutes and 32 seconds.

### Data pre-processing

The Elekta Neuromag MaxFilter 2.2 has been applied to all MEG data (temporal signal space separation (tSSS): 0.98 correlation, 10s window; bad channel correction: ON; motion correction: OFF; 50Hz+harmonics (mains) notch). Afterwards, artifact rejection, filtering (0.3-100 Hz bandpass), re-referencing (i.e. using the algebraic average of the left and right mastoid electrodes), temporal segmentation into epochs, averaging and source estimation were performed using Brainstorm (Tadel *et al*., 2011). In addition, physiological artefacts (e.g. blinks, saccades) were identified and removed using spatial space projection of the signal. In order to improve the accuracy of the source reconstruction, the FreeSurfer (Fischl, 2012), software was used to generate cortical surfaces and automatically segment them from the cortical structures from each participant’s T1-weighted anatomical MRI. The advanced MEG model was obtained from a symmetric boundary element method (BEM model; OpenMEEG; Gramfort *et al*., 2010; Kybic *et al*., 2005), fitted to the spatial positions of each electrode (Huang *et al*., 1999). A cortically constrained sLORETA procedure was applied to estimate the cortical origin of the scalp MEG signals. The estimated sources were then smoothed and projected into a standard space (i.e., the ICBM152 model) for comparisons between groups and individuals, while accounting for differences in native anatomy. This procedure was applied for the entire recording duration.

### Network segmentation

In line with previous work (e.g., Smitha *et al*., 2017; Van den Heuvel *et al*., 2009), we investigated the four main brain networks at rest: the default-mode network (DMN), the salience network (SN), the left fronto-parietal network (FPL) and the right fronto-parietal network (FPR). Each network is composed of different brain regions: the DMN is composed of the posterior cingulate cortex, the medial prefrontal and the inferior parietal cortex. The SN is composed of the anterior cingulate cortex, the insula and the pre-supplementary motor area. The FPL is composed of the left dorsolateral prefrontal cortex and the left superior parietal cortex. Finally, the FPR is composed of the right dorsolateral prefrontal cortex and the right superior parietal cortex (Figure 1). These networks are involved in different cognitive activities or functions: the DMN is mainly observed at rest and shows lower connectivity levels when participants are currently performing cognitive tasks (Raichle *et al*., 2001). The SN is associated with the processing of salient stimuli in the environment (Seeley *et al*., 2007). Finally, the bilateral fronto-parietal network is involved in spatial attention, planning and cognitive control (Kam *et al*., 2019). We separately investigated the FPL, which is involved in working memory (Murphy *et al*., 2019) and the FPR which is involved in inhibitory processing (Nee *et al*., 2007). Regions of interest were selected following segmentation of individual anatomies based on the Desikan-Killiany atlas (Desikan *et al*., 2006).

**Figure 1:**
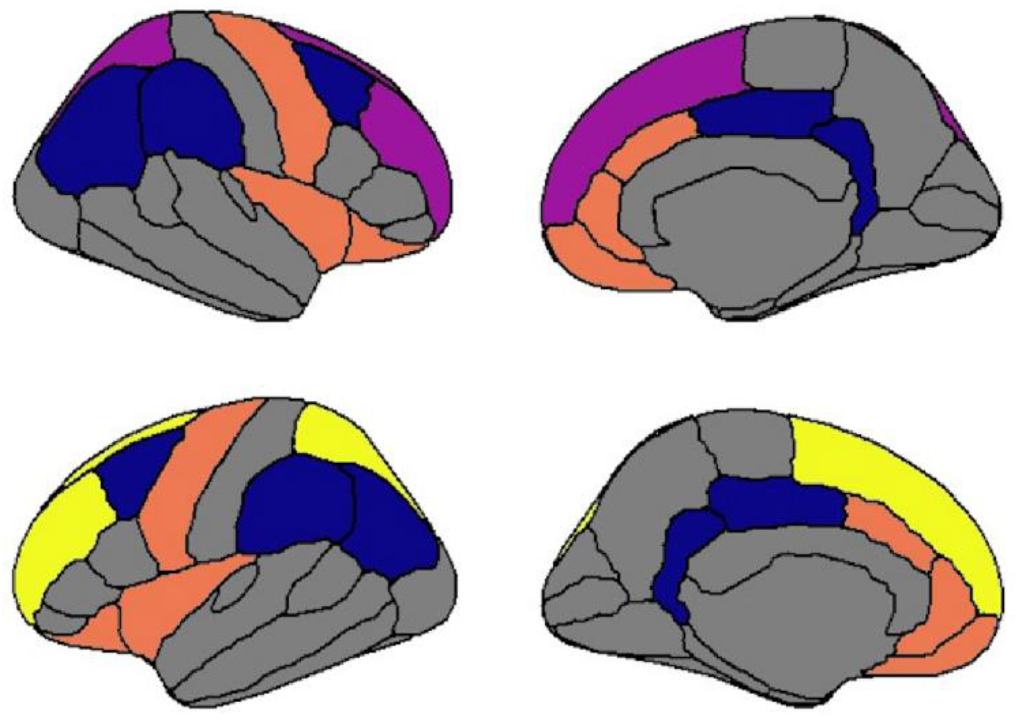
Visualisation of the different regions forming the four studied brain networks;in blue: the DMN; in orange: the SN; in yellow: the FPL; in purple: the FPR

### Study of dynamic connectivity

Phase-locking value analyses (PLV; Lachaux *et al*., 1999) were used to determine the functional synchrony between regions of interest. PLV estimates the variability of phase differences between two regions over time. If the phase difference varies little, the PLV is close to 1 (this corresponds to high synchronisation between the regions), while the low association of phase difference across regions is indicated by a PLV value close to zero. To ensure PLV results did not reflect volume conduction artefacts, control analyses were conducted using phase lag index (weighted PLI analyses). Because PLV is an undirected measure of functional connectivity, and to investigate brain dynamics with complementary metrics, analyses of transfer entropy (TE) have also been conducted. TE measures of how a signal a can predict subsequent changes in a signal b (Ursino *et al*., 2020). It then provides a directed measure of a coupling’s strength. If there is no coupling between a and b, then TE is close to 0, while TE is close to 1 if there is a strong coupling between a and b.

The range of each frequency band was based on the frequency of the individually observed alpha peak frequency (IAF), measured as the average of peaks detected with both occipitoparietal magnetometers and gradiometers. From previous work (Toppi *et al*., 2018) the following frequency bands were considered: Delta (IAF-8/IAF-6), Theta (IAF-6/IAF-2), Alpha (IAF-2/IAF+2), Beta (IAF+2/IAF+14), Gamma1 (IAF+15/IAF+30) and Gamma2 (IAF+31/IAF+80). To reduce the dimensionality of the data, the first principal component analysis (PCA) decomposition mode of the time course of activation in each region of interest (ROI) of the Desikan-Killiany atlas brain fragmentation was used. The first component, rather than the average activity, was chosen to reduce signal leakage (Sato *et al*., 2018). 35 sliding 30s sliding time windows were then extracted for the epochs of interest to calculate the variability across time windows (standard deviation) of the PLV. The analyses were conducted on the average activity within each network, however additional analyses were conducted at the coupling level to further investigate the observed results.

### Statistical tests

Permutation analyses were performed in Brainstorm (Tadel et al., 2011), using methods originally implemented in Fieldtrip (Maris & Oostenveld et al., 2011). Both toolboxes support open access and scripts are available online. To assess differences between age groups in demographic and functional connectivity variables, t-tests and ANOVAs were applied using Jamovi software (https://www.jamovi.org/; version 1.6.23). Functional data (PLV, TE) were analyzed using 2 (age group: young/old) x 4 (networks: DMN, SN, FPL, and FPR) x 6 (frequency bands: delta, theta, alpha, beta, gamma1, gamma2) repeated-measures ANOVAs to determine which network and frequency band showed the greatest young/old changes. The Greenhouse-Geisser epsilon correction was used where necessary. Original degrees of freedom and corrected p-values are reported. Finally, regressions aimed at determining the association between functional connectivity measures and behavioral measures within each group. Results were FDR corrected for multiple comparisons (Benjamini & Hochberg, 1995).

## Results

### Age-related differences in cognitive performance

The main behavioral and demographic data from the Cam-CAN database are summarized in Table 1.

Relative to younger individuals, older adults showed lower scores in the MMSE (p=0.013), VSTM (p<0.001), Cattell (p<0.001) and hotel test (p=0.018 for number of rooms; and p=0.005 for time) scores. For the hotel test, a decrease in the rate of correct answers was observed (p=0.018). A significant increase in response time for the hotel test was also observed in older individuals (p=0.005).

### Increased variability of delta phase synchrony frequency band in older adults

We first observed a significant effect of network, F(3, 270) = 8.085, p<0.001, η^2^ = 0.082, frequency, F(5, 450) = 202.748, p<0.001, η^2^ = 0.693, and age, F(1,90) = 4.698, p= 0.033, η2= 0.05. The interaction between frequency and age, F(5,450) = 6.57, p<0.001, η^2^ = 0.068, revealed that this difference in variability between young (M = 0.076, SE = 0.002) and older adults (M = 0.087, SE = 0.002) was stronger for the delta frequency band. This effect was not observed in other frequency bands. The Age x Networks interaction for the delta frequency band was also significant, F(3,270) = 6.823, p<0.001, η^2^ = 0.07, with the DMN network showing the largest difference. These results indicate an increased variability of the delta DMN activity with advancing age (Figure 2). We observed a significant negative regression between such variability and cognitive performance (VSTM, p = 0.009, r = −0.387). The rest of the analyses was therefore focused on the DMN network, in the delta frequency band.

**Figure 2:**
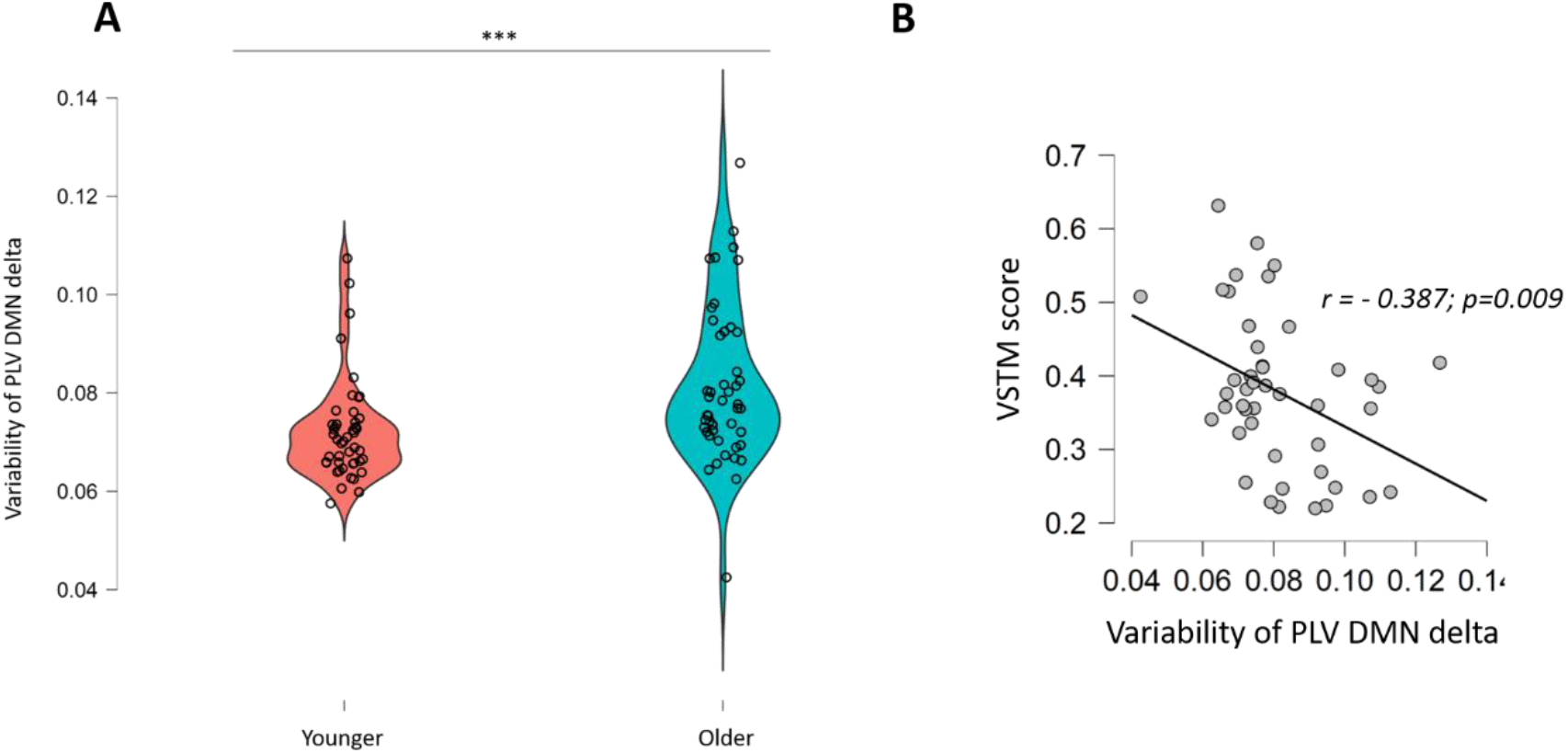
**A**: Increased variability of the DMN network in the delta frequency band for the older group (p = 0.001) compared to younger individuals; **B**: Negative association between increased PLV DMN variability and VSTM score (regression test, r = −0.387, p = 0.009) in older adults

We then performed permutation t-tests on the DMN couplings between age groups. Different couplings were found to be significantly more variable for the older group compared to the younger group especially for interhemispheric and fronto-parietal couplings (Figure 3). In the older group, the right frontoparietal coupling was found to be negatively correlated with cognitive performance (MMSE test; r = - 0.305, p = 0.039). In the older group, the interhemispheric coupling (bilateral supramarginal regions) was found to be negatively correlated with cognitive performance (VSTM test; r = −0.344, p = 0.021). These data suggest an increase in variability in the overall DMN network in the delta frequency band, but also an increase in the significant variability of specific couplings in this network, both being associated with lower cognitive performance.

**Figure 3:**
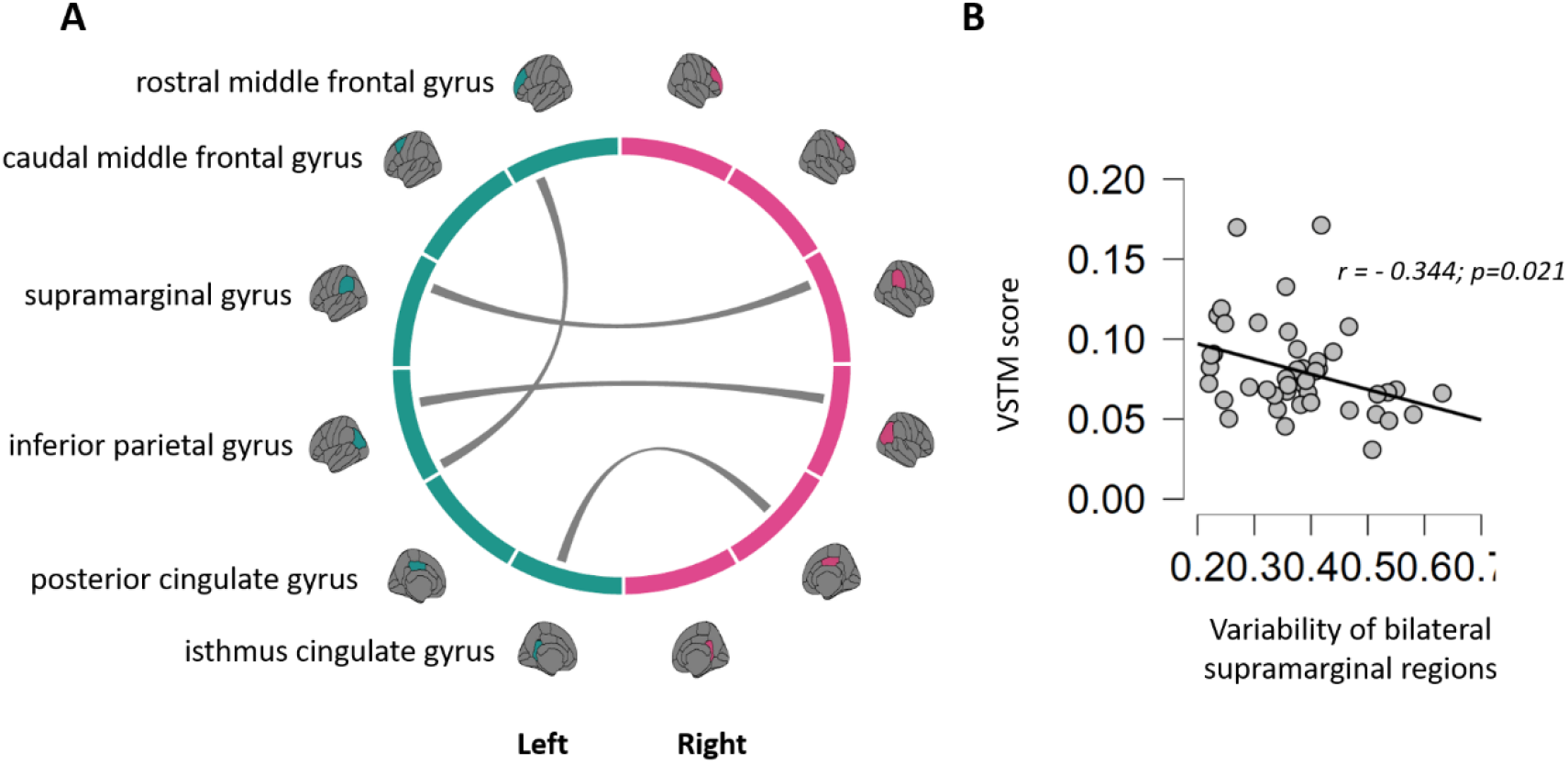
**A**: Increased PLV variability of the DMN couplings in the delta frequency band for the older group (p = 0.001); **B**: Negative association between increased variability of PLV DMN coupling (bilateral supramarginal regions) and Cattell score (regression test, r =-0.344, p = 0.021) in older adults

### Reversal of the direction of information transfer of delta band in older adults

As phase synchrony measures are undirected, transfer entropy was used to determine whether a specific direction of connectivity was associated with age-related differences. We performed a repeated measures ANOVA (Age x Networks x Frequencies x Direction) to determine which, network, which frequency band and in which direction the largest young-to-old changes were found. We showed a significant effect of frequency, F(5, 450) = 361.1, p<0.001, η^2^ = 0.801. A significant effect of age, F(1, 90) = 17.7 p<0.001, η^2^= 0.165 was also observed. Results revealed an increase in the direction of information transfer variability in the delta frequency band, in older adults relative to young adults. An interaction between frequency and age was also observed, F(5, 450) = 14.61, p<0.001, η^2^ = 0.140. This significant interaction effect indicates larger coupling strength in delta frequency in the older group (M = 1.218, SE = 0.0231) compared to the younger group (M = 0.921, SE = 0.0231).

Student’s t-tests were performed to determine the direction of information transfer for young and older adults in the DMN. We saw a significant difference between the fronto-parietal and parieto-frontal direction (p=0.013), with a significantly larger coupling strength parieto-frontal direction for young relative to older participants. With advancing age, in the DMN network and the delta frequency band, a decrease in the transfer of information from parietal to frontal regions has been observed. This decrease in communication can be linked to the cognitive performance observed in this group. Indeed, we conducted regressions analyses to determine the association between these entropy transfer measures and behavioural measures within each group. We found negative regressions of information transfer with cognitive performance in all directions (VSTM, p=0.031, r= −0.319; Cattell, p=0.020, r= −0.341) in the delta frequency band for older adults (Figure 4).

**Figure 4:**
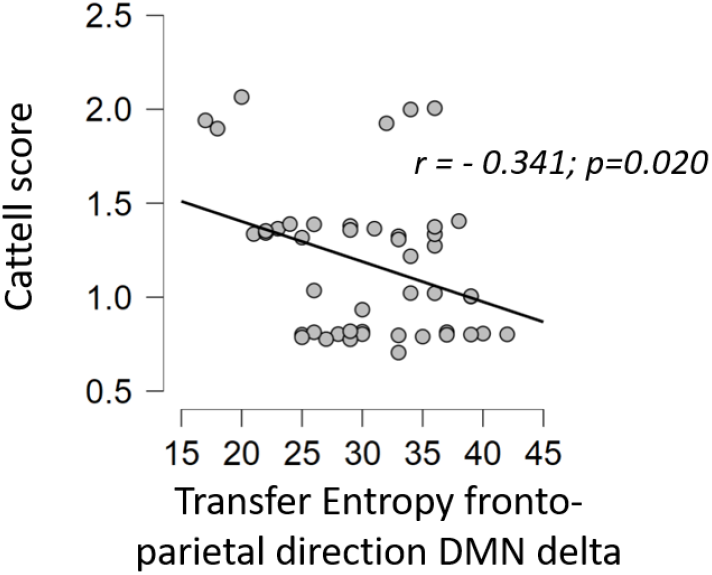
Negative association between increased of parieto-frontal direction in DMN network and Cattell score (regression test, r = - 0.341, p = 0.020)

## Discussion

Our main objective was to investigate changes in the stability and variability of brain communication dynamics with age and the relationship of these changes with age-related cognitive changes. Our connectome-based approach, based on MEG data in healthy young and older participants from the Cam-CAN database, allowed us to investigate changes of connectivity dynamics with aging. Two time-resolved connectivity aspects were studied: the stability of synchronized communications over time, and directed connectivity. Brain activity was studied at rest, as previous work suggested a link between the activity of specific networks at rest and cognitive abilities (e.g. Nashiro *et al*., 2017). In this study, we first showed an increased variability of phase synchrony over time with age, especially in the delta frequency band. We also showed a reversal of the main direction of synchronized connectivity with age: connectivity in the fronto-parietal direction was found to be increased in older participants, whereas it was stronger in the parieto-frontal direction for younger participants. These observations are in line with the available literature on functional connectivity during non-pathological aging (e.g. Geerligs *et al*., 2015). The results also show for the first time that the stability or variability of functional networks, as well as information exchange over time, are associated with individual cognitive differences during aging. This was made possible by the excellent temporal resolution of MEG, combined with advanced source reconstruction analyses.

The study of oscillatory activity allowed us to specify age-related changes in the variability of phase synchrony over time, and the specific frequency band associated with these differences. Phase synchrony between brain regions is a critical parameter of neural communications (e.g., Fries, 2015). Indeed, with advancing age, changes in synchronized network communications have been observed (see Courtney & Hinault, 2021, for a review). Our results reveal an increased variability of phase synchrony in the default network, mainly in the delta frequency band with age. Such variability of neural synchrony was negatively correlated with cognitive performance (measures of general cognition, and working memory). This result is consistent with MRI work showing that an age-related decrease in connectivity within the DMN is related to a decrease in memory and executive functions (e.g. Andrews-Hanna *et al*., 2007). Our results are also consistent with previous M/EEG work reporting an overall slowing of brain activity with advancing age (e.g., Celesia, 1986), with an increase of slow rhythms relative to faster rhythms. Increased slow waves seem to be associated with the cognitive decline observed with advancing age. Here, we show that this slowing of brain rhythms with age is associated with a loss of stability in neuronal communications, and poorer performance.

In association with synchrony analyses, transfer entropy analyses allow the quantification of directed connectivity (see Ursino *et al*., 2020). This quantifies the information flow between brain regions more precisely than functional connectivity, thus allowing the detection of causal interactions (i.e., *A* must precede *B*) between brain regions. Such investigation of directed connectivity revealed a decrease in the parieto-frontal direction of brain communications relative to the fronto-parietal direction in the default network and the delta frequency band with age. This reversal of information transfer between young and old participants was negatively correlated with cognitive performance (especially for working memory and fluid intelligence). The reversal of information transfer and decreased variability in phase synchrony observed here may help furthering the age-related pattern described in the PASA model (Cabeza *et al*., 2018). According to this model, the increase in the recruitment of frontal regions in older adults would be an indicator of their attempts to compensate for the decrease in their cognitive abilities. Here, we show a decrease in information transfer to these frontal regions, which is negatively associated with cognitive performances. Reduced connectivity of these frontal regions has been found to be negatively correlated with cognitive performance (Toth et al., 2014), which may reflect a decrease in recruitment to these frontal regions. Indeed, frontal regions are the first regions to see their neuroanatomy impacted by aging (Dennis & Cabeza, 2008). These results allow us to understand this concept at the network communication dynamics levels. Further investigation of investigation transfer during task completion will be necessary to specify its associations with the direct implementation of cognitive processes.

Several methodological considerations must be discussed regarding the reported results. First, the investigation of resting-state activity prevents in part the direct investigation of the neural bases of cognitive processes, which may explain the small number of associations with cognition. This could also reflect the fact that the Cam-CAN database does not include tasks directly testing executive functions. However, studying dynamic network connectivity at rest furthers our knowledge on the stability of these networks and help better characterize their individual variations. Second, the use of the Desikan-Killiany atlas, which has a less precise spatial resolution than other atlases, could limit the interpretations of SN results. However, the spatial resolution of the MEG does not enable a much higher spatial resolution. The Desikan atlas allows to limit the degrees of freedom and has been frequently reported in previous work (e.g., Ceisnaite et al., 2021; Canal-Garcia et al., 2022). Third, eta-squares for the effect of age on variance and transfer entropy show small to medium effects. Nevertheless, observed these effects in healthy older adults, at rest, suggest they are sensitive to early age-related changes. Future work is necessary to assess longitudinal changes of dynamic connectivity. Finally, the use of PLV for M/EEG data can be problematic because of volume conduction effects. However, volume conduction effects are unlikely to explain couplings between distant brain regions, and cannot account for between-group differences or correlations with behavioral performance. Using the first mode of principal component analysis (PCA) of the activation time course in each region of interest, rather than mean activity, reduces signal leakage (e.g., Sato et al 2018). Finally, even alternative measures (such as Phase Lag Index (PLI), or coherence), do not fully control for this risk of conduction volume (Palva et al 2017). Additional weighted PLI analyses were conducted, and results were replicated for the DMN network in the delta frequency band.

Theoretically, the variability of brain communications has received little investigation, as it was long considered as noise, but is now recognized as contributing to brain functions (Uddin *et al*., 2020). Here, we show that healthy aging is associated with an increased variability in synchronized brain communications, and with changes of the main connectivity directions between brain regions. Results highlight that even when brain networks are not engaged in a particular cognitive activity, significant changes occur with age regarding connectivity dynamics and information flow between regions of different functional brain networks. Advancing age appears to be accompanied by a functional disorganization of dynamic networks, with a loss of communication stability and a decrease in the information transmitted. The study of dynamic connectivity contributes to a better understanding of the cognitive decline with aging. The stability of communications and its alteration should be considered in the framework of maintenance, reserve and resilience (Cabeza 2018; Stern, 2020).

## Acknowledgments

This research did not receive any specific grant from funding agencies in the public, commercial, or not-for-profit sectors.

## Notes

**Declarations of interest**: none

### Competing Interest Statement

The authors have declared no competing interest.

